# Photoreceptor progenitor dynamics in the zebrafish embryo retina and its modulation by primary cilia and N-cadherin

**DOI:** 10.1101/2020.02.13.947663

**Authors:** Gonzalo Aparicio, Magela Rodao, José L. Badano, Flavio R. Zolessi

**Affiliations:** Sección Biología Celular, Facultad de Ciencias, Universidad de la República, Uruguay; Institut Pasteur Montevideo, Uruguay

**Author notes:** contributed equally to this work (Gonzalo Aparicio, Magela Rodao). ^†^Corresponding Author: Flavio R. Zolessi, Sección Biología Celular, Facultad de Ciencias, Universidad de la República., Calle Iguá 4225, Montevideo 11400, Uruguay. Co-author emails: GA,; MR,; JLB.

**Keywords:** neuroepithelium, cell polarity, cell adhesion, retina lamination, neurogenesis

## Abstract

**Background:** Photoreceptors of the vertebrate neural retina are originated from the neuroepithelium, and like other neurons, must undergo cell body translocation and polarity transitions to acquire their final functional morphology, which includes features of neuronal and epithelial cells.

**Methods:** We analyzed this process in detail on zebrafish embryos using *in vivo* confocal microscopy and electron microscopy. Photoreceptor progenitors were labeled by the transgenic expression of EGFP under the regulation of the photoreceptor-specific promoter *crx*, and genes of interest were knocked-down using morpholino oligomers.

**Results:** Photoreceptor progenitors detached from the basal retina at pre-mitotic stages, rapidly retracting a short basal process as the cell body translocated apically. They remained at an apical position indefinitely to form the outer nuclear layer (ONL), initially extending and retracting highly dynamic neurite-like processes, tangential to the apical surface. Many photoreceptor progenitors presented a short apical primary cilium. The number and length of these cilia was gradually reduced until nearly disappearing around 60 hpf. Their disruption by knocking-down IFT88 and Elipsa caused a notorious defect on basal process retraction. Time-lapse analysis of N-cadherin knock-down, a treatment known to cause a severe disruption of the ONL, showed that the ectopic photoreceptor progenitors initially migrated in an apparent random manner, profusely extending cell processes, until they encountered other cells to establish cell rosettes in which they stayed acquiring the photoreceptor-like polarity.

**Conclusion:** Altogether, our observations indicate a complex regulation of photoreceptor progenitor dynamics to form the retinal ONL, previous to the post-mitotic maturation stages.

## Introduction

The central nervous system of vertebrates is originated from a polarized, pseudostratified epithelium, which among other features presents a Laminin1-enriched basal lamina, a sub-apical N-cadherin-based adherens junctions and apical primary cilia (Zolessi, 2016). These neuroepithelial cells are the progenitors for both glial cells and neurons. It is hence natural to assume that the epithelial polarized surroundings must influence how neurons acquire their own polarity, and, particularly, orientation inside the tissue. The identity of these signals and their interactions with the early post-mitotic neurons are, however, still poorly understood. The neural retina is a highly approachable region of the central nervous system, ideal for i*n vivo* studies of neural differentiation (Amini *et al.*, 2018). In spite of its relative simplicity, it contains a diversity of cell types, ranging from the very “canonical” neurons such as retinal ganglion cells (RGCs; with a single axon, and an opposite branching dendrites), to neurons that partly remain as epithelial cells, like the photoreceptors (Hoon *et al.*, 2014). Mutant analyses in zebrafish embryos have indicated that the histogenesis of the retina appears particularly sensitive to the disruption of molecules involved in epithelial cell polarity. Most of the affected genes in these mutants encode for proteins related to apical identity and sub-apical adhesion in epithelial cells, such as N-cadherin (mutants known as *parachute, glass onion* or *lyra*), Pals1/Mpp5 (*nagie oko*) and aPKCλ (*heart and soul*) and Crb2a (*oko meduzy*) (Pujic and Malicki, 2004).

Differentiated photoreceptors, cones and rods, are unique cells in many respects. First, the largest part of their cytoplasm and membrane protrudes from the apical side of the cell, beyond the adhesion belt that holds them together with the apical tips of Müller glia (the “outer limiting membrane”). This apical portion consists of the inner segment, highly enriched in membrane organelles and mitochondria, and the outer segment, where a thick stack of membranes contains the photo-transduction machinery. Interestingly, the outer segment is originated from and remains attached to the inner segment through a connecting cilium, essential for the transport of proteins and membrane (Malicki, 2012). The photoreceptor cell body and nucleus are mostly located below the outer limiting membrane, and basally the cell ends in a specialized pre-synaptic terminal, connecting with bipolar and horizontal cells. Alterations in the maintenance of this particular cell polarity, as happens when genes encoding proteins involved in apical adhesions or cilia transport are mutated, can eventually cause retinal degeneration in animals and humans. For example, mutations in Crb genes cause hereditary forms of Leber congenital amaurosis and retinitis pigmentosa (Quinn *et al.*, 2017), while a common symptom in ciliopathies is retinal dystrophy (Wheway *et al.*, 2014). Studying how photoreceptors differentiate *in vivo*, particularly at earlier developmental stages when the first polarized signals act on progenitors and early post-mitotic cells, is of vital importance for understanding how these diseases eventually arise.

By *in vivo* imaging, it was shown that photoreceptors arise from terminal cell divisions of apically-localized progenitors that already display an epithelial-like conformation, reminiscent of the organization at the mature outer nuclear layer, indicating that polarity constraints are acting very early on the photoreceptor cell lineage (Suzuki *et al.*, 2013; Weber *et al.*, 2014). This process appears to be severely affected in all polarity mutants described above. For example, in *nagie oko* mutant retinas it was shown that photoreceptors have a scattered distribution, while in different N-cadherin mutants they tend to form rosettes inside the retina (Masai *et al.*, 2003; Wei *et al.*, 2006). This effect, as it was shown for similar mutants, was non-cell autonomous, indicating the importance of cell-to-cell interactions and adherens junctions during the formation of the outer nuclear layer (ONL; Wei *et al.*, 2006; Zou *et al.*, 2008). In this work, we provide a detailed description of the normal dynamics at early stages of photoreceptor cell differentiation, starting at progenitor stages, as labeled by using fluorescent reporters under the regulation of the *crx* promoter region. In addition, we analyzed by confocal and electron microscopy the presence and localization of primary cilia in these precursor cells, well before the onset of outer segment formation. Finally, we disrupted cilia, as well as adhesion complexes mediated by N-cadherin, using well-characterized morpholinos (Lele *et al.*, 2002; Lepanto, Davison, *et al.*, 2016). Our observations indicate different roles for primary cilia and N-cadherin on photoreceptor precursor organization in the intact zebrafish retina, with only the latter appearing essential for cell orientation.

## METHODS

### Fish care and breeding

Zebrafish were maintained and bred using standard methods (Lepanto et al., 2016b). Embryos were raised at 25 to 31 °C depending on the experiments, and staged in hours post-fertilization (hpf). We used wild-type (Tab5) and previously established transgenic lines: Tg(*actb2*:Arl13b-EGFP)hsc5 (Arl13b-GFP), kindly provided by B. Ciruna (Borovina *et al.*, 2010); Tg(*atoh7*:gap43-mRFP)cu2 (Zolessi *et al.*, 2006). All manipulations were carried out following the approved local regulations (CEUA-IPMon, and CNEA).

### Injection of molecular constructs and morpholino oligomers (MOs)

We constructed the following vectors using the Multisite Gateway-based (ThermoFisher Scientific) tol2 kit system (Kwan *et al.*, 2007): pDestTol2pA2;*crx*:EGFP-CAAX, pDestTol2pA2;*crx*:mCherry-CAAX. p5E-*crx* was kindly provided by R. Wong (Suzuki *et al.*, 2013). The construct *bact2*:APEX2-GBP cloned in a pDESTTol2pA2 was obtained from Addgene (Plasmid 67668) (Ariotti *et al.*, 2015). Plasmid DNA together with Tol2 transposase mRNA (0.5 nL; 2.5 and 6 pg, respectively) were injected at the one-cell stage according to standard techniques. For transgenic line generation, identified carriers were outcrossed until offspring inherited the transgene at mendelian ratios. We generated the following transgenic lines in this work: Tg (*crx*:EGFP-CAAX,*cmlc2*:GFP); Tg (*crx*:mApple-CAAX, *cmlc2*:GFP); Tg(*bact2*:APEX2-GBP,*acrys*:mCherry).

Previously well-characterized MOs were obtained from Gene Tools (Philomath, USA): ift88-SP (AACAGCAGATGCAAAATGACTCACT), which targets the exon 3 - intron 3 boundary of *ift88* (Lepanto, Davison, *et al.*, 2016); *elipsa*-SP (CTGTTTTAATAACTCACCTCGCTGA) which targets the exon 1 - intron 1 boundary of *elipsa* (Lepanto, Davison, *et al.*, 2016); *cdh2*-TR (TCTGTATAAAGAAACCGATAGAGTT) which targets the 5’ UTR of the N-cadherin mRNA (Lele *et al.*, 2002). All MOs were injected in the yolk of 1–4 cell-stage embryos, at a maximum volume of 3 nL. As a control, we used matching doses of a standard MO (CCTCTTACCTCAGTTACAATTTATA) from Gene Tools (Philomath, USA). In all cases, we co-injected the standard anti-p53 MO (Robu *et al.*, 2007).

### Immunofluorescence

Embryos grown in 0.003 % phenylthiourea (PTU; Sigma) were fixed overnight at 4 °C in in 4 % paraformaldehyde in PBS and subjected to permeabilization and whole-mount immunostaining as previously described (Lepanto et al., 2016b). Primary antibodies used: anti-acetylated tubulin (Sigma-Aldrich, T-7451), 1/1000; anti-γ-tubulin (Sigma-Aldrich, T6557), 1/500; anti-GFP (DSHB, DSHB-GFP-12A6), 1/200; anti-pan-cadherin (Sigma-Aldrich, C3678), 1/500; anti-aPKCζ (SCBT, sc-216), 1/500. Secondary antibodies from ThermoFisher Scientific: anti-mouse IgG-Alexa 488 (A11034), 1/1000; anti-mouse IgG-Alexa 555 (A21424), 1/1000; anti-rabbit IgG-Alexa 633 (A21070). All antibody incubations were performed overnight at 4 °C. Nuclei were fluorescently labeled using methyl green (Prieto et al, 2014) and actin filaments with TRITC-conjugated phalloidin (Sigma-Aldrich, P1951). Observation of whole embryos was performed using a Zeiss LSM 880 or a Zeiss LSM 800 laser confocal microscope, with 20x 0.7 NA, 25x 0.8 NA, 40x 1.2 NA or 63x 1.3 NA glycerol:water (75:25).

### In vivo confocal microscopy

36 hpf embryos were selected, anesthetized using 0.04 mg/mL MS222 (Sigma) and mounted in 1 % low melting-point agarose, containing 0.003 % N-phenylthiourea and 0.04 mg/ml MS222/tricaine (Sigma) over n° 0 glass bottom dishes (MaTek). During overnight image acquisitions, embryos were kept in Ringer’s solution (116 mM NaCl, 2.9 mM KCl, 1.8 mM CaCl2, 5 mM HEPES pH 7.2) with 0.04 mg/mL MS222. Live acquisitions were made using a Zeiss LSM 880 laser confocal microscope with a 25X 0.8 NA or 40x 1.2 NA objective and glycerol:water (75:25) immersion medium. Stacks around 50 μm-thick were acquired in bidirectional mode, at 1 μm spacing and 512 × 512 pixel resolution every 10 or 15 min. The acquisition time per embryo was approximately 45 s, and up to 8 embryos were imaged in each experiment.

### Transmission electron microscopy

Embryos were fixed for 30 min at RT by immersion in 4 % paraformaldehyde, 2.5 % glutaraldehyde, 0.1 M phosphate buffer, pH 7.2–7.4. Heads were then dissected and incubated in fixative solution overnight at 4 °C. After washing, they were post-fixed in 1 % osmium tetroxide. APEX2-GBP embryos were processed according to Ariotti *et al.*, 2015. Briefly, embryos were fixed for 30 min at room temperature by immersion in 4 % paraformaldehyde, 2.5 % glutaraldehyde, 0.1 M sodium cacodylate buffer, pH 7.4. Dissected heads were then fixed overnight at 4 °C. After repeatedly washing in buffer, embryo heads were incubated with diaminobencidine (DAB) in the presence of H2O2 for 30 min at room temperature, washed in 0.1 M sodium cacodylate buffer, post-fixed in 1% osmium tetroxide for 15 min and washed again. In all cases, embryo heads were dehydrated using ethanol, infiltrated and embedded in Araldite resin. Ultrathin 70 nm sections were obtained using an RMC MT-X ultramicrotome, mounted on formvar-coated copper grids and stained with 2 % aqueous uranyl acetate followed by Reynold’s lead citrate. Observation and acquisition was performed using a Jeol JEM 1010 transmission electron microscope operated at 80 kV, equipped with a Hamamatsu C4742-95 digital camera or a Jeol JEM 2100 operated at 120 kV, with a Gatan Orius 1000 digital camera.

### Data analysis

Images were processed and analyzed using Fiji (https://fiji.sc/). Statistical analyses were performed using GraphPad Prism 6. As a routine, the datasets were checked for normality using the D’Agostino-Pearson, or the Shapiro-Wilk normality tests. To analyze statistical significance of differences in average, we performed a Student’s *t* test in the case of normal data distribution, and a Mann–Whitney test in the case of non-normal data.

## RESULTS

### Dynamic behavior of photoreceptor progenitors during the establishment of the outer nuclear layer (ONL)

We labeled photoreceptors using the regulatory region of the cone-rod homeobox gene (*crx*) to drive the expression of membrane-directed EGFP-CAAX (*crx*:GFP), a construct that is expressed from progenitor stages (Suzuki *et al.*, 2013). Similar to what has been previously shown (Weber *et al.*, 2014), we observed here a gradual formation of the ONL, between approximately 36 and 60 hpf (Fig. 1A). This is evidenced by transmission electron microscopy (TEM) as well (Fig. 1B), where we could detect photoreceptor progenitors at early stages like 36 hpf by crossing *crx*:GFP transgenic with APEX-GBP transgenic lines (Ariotti *et al.*, 2015). At 48 hpf, the photoreceptor progenitors already formed a highly ordered sheet of cells at the apical side of the retina, while cell divisions were still evident (Fig 1). The epithelial-like organization of the ONL at 48 and 60 hpf was confirmed by immunolabeling with anti-pan-cadherin and anti-aPKCζ antibodies, showing an apical accumulation of both molecular markers at these stages on *crx*:GFP-expressing cells (Fig. 1D,E). Interestingly, F-actin appeared to be accumulated at a thinner apical region of these cells at 48 than at 60 hpf, with a relatively higher coincidence with cadherins at the later stage (Fig. 1D). At 48 hpf, most of the cadherin signal was excluded from the thinner apical F-actin-rich zone. In the case of the apical marker aPKCζ, however, signal was always restricted to the apical-most aspect of *crx*:GFP cells, apparently never coinciding with the basal extension of the F-actin accumulation, where the pan-cadherin signal was evident at 60 hpf (Fig. 1E).

**Figure 1.**
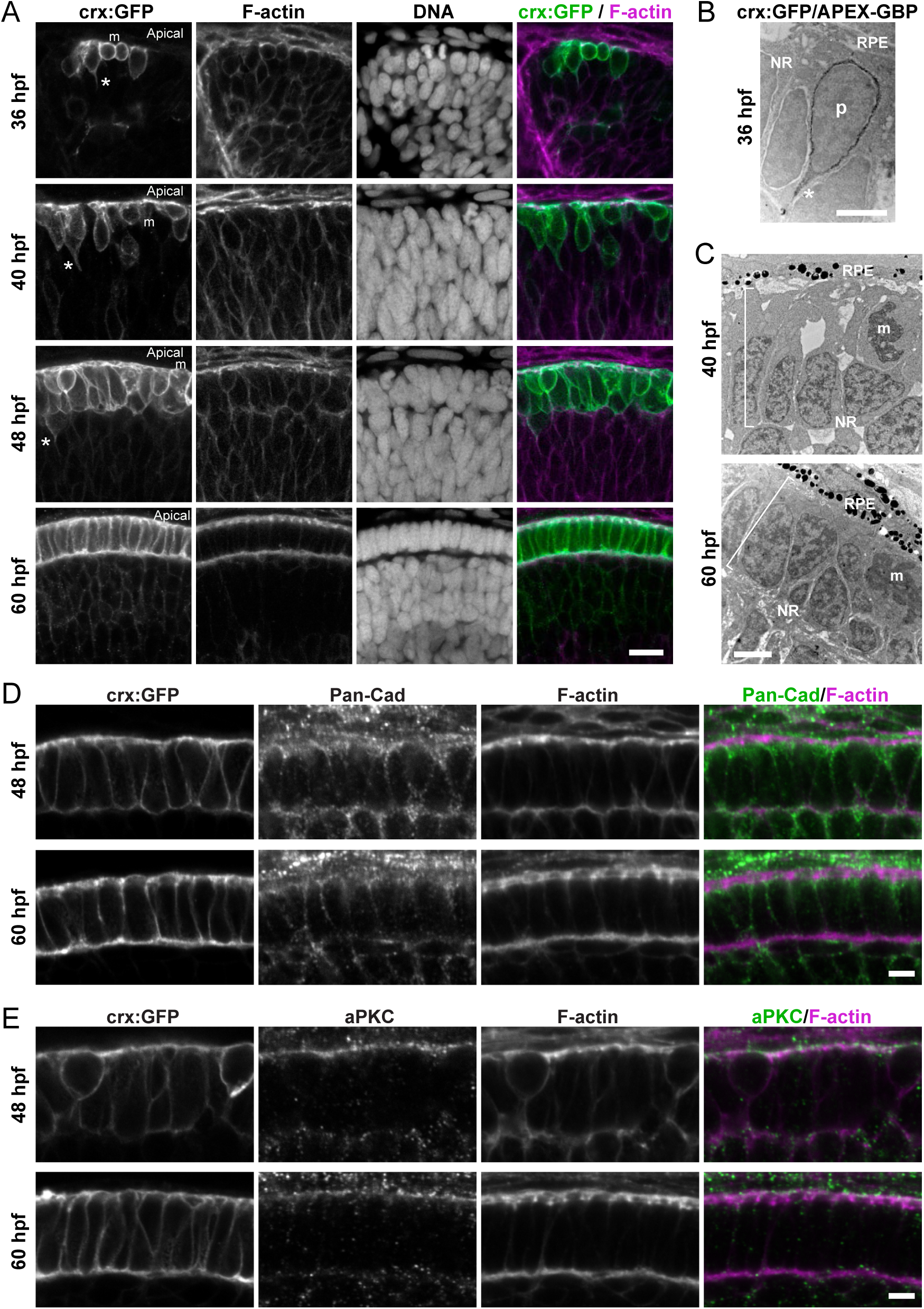
Formation of the outer nuclear layer in the early stages of retina histogenesis. **(A)** Series of confocal images from zebrafish embryo retinas at different developmental stages, transgenically labeled with *crx*:EGFP-CAAX (*crx*:GFP). m: mitotic cells; asterisks: basal processes. 36 and 60 hpf, single confocal sections; 40 and 48 hpf, maximum intensity projections of four sections, at 0.37 µm separation. **(B)** Transmission electron microscopy (TEM) of *crx*:GFP/APEX-GBP double transgenic embryo at 36 hpf. Binding of the modified peroxidase APEX-GBP to GFP allows for the generation of a DAB precipitate around the cell periphery of photoreceptor progenitors. NR: neural retina; RPE: retinal pigmentary epithelium; asterisk: basal process. **(C)** TEM of the outer retina at later stages; the ONL (brackets) starts to become evident as photoreceptor progenitors elongate and acquire an epithelial-like conformation. NR: neural retina; RPE: retinal pigmentary epithelium; m: cells undergoing mitosis. **(D-E)** High magnification confocal images of the apical region of *crx*:GFP-transgenic embryos labeled with anti-pan-cadherin antibody (D) and anti-aPKCζ antibody (E). Scale bars: A, 10 µm; B - C, 2 µm; D-E: 5 µm.

Time-lapse confocal microscopy of *crx*:GFP-expressing cells (from 36 hpf) showed a progressive signal increase, with a relatively good detection of photoreceptor progenitors localized around the central area of the retinal neuroepithelium, from where they translocated their nuclei apically, possibly during G2 phase of the cell cycle (Fig. 2A,B and Supplementary Video 1). Although difficult to detect in general, we could observe in a few cases an onset of *crx*:GFP expression in cells that appeared to be attached to the basal side of the neuroepithelium through a very thin basal process, which was quickly retracted by the time of apical nuclear translocation (Fig. 2C and Supplementary Video 2). While their nuclei moved apically, some photoreceptor progenitors presented a short basal process, while others just had a smooth basal side (Fig. 1A and 2A,B). All along the process, the observed cells remained attached to the apical border of the retinal neuroepithelium, initially through an apical process that quickly shortened until it disappeared as the cell body translocated to the future ONL (Fig. 2A,B and Supplementary Video 1). This process of nuclear translocation was clearly seen in most *crx*:GFP-expressing cells at 36 hpf, and in cells located at the periphery of the central patch of differentiating retina at 40 and 48 hpf (also see Fig. 3 and associated Supplementary Videos). A quantification of cell body translocation showed that since clearly detected, photoreceptor progenitors presented a net directional movement towards the apical border of the neuroepithelium, but with varying speeds and eventual reversed direction, taking a total time between 2 and 4 hours (Fig. 2D,E). Once translocated, photoreceptor progenitors apparently stayed permanently in that apical position, eventually dividing between 5 and 8 h after arriving to the ONL without any visible interkinetic nuclear migration (Fig. 2F,G and Supplementary videos 1 and 3).

**Figure 2.**
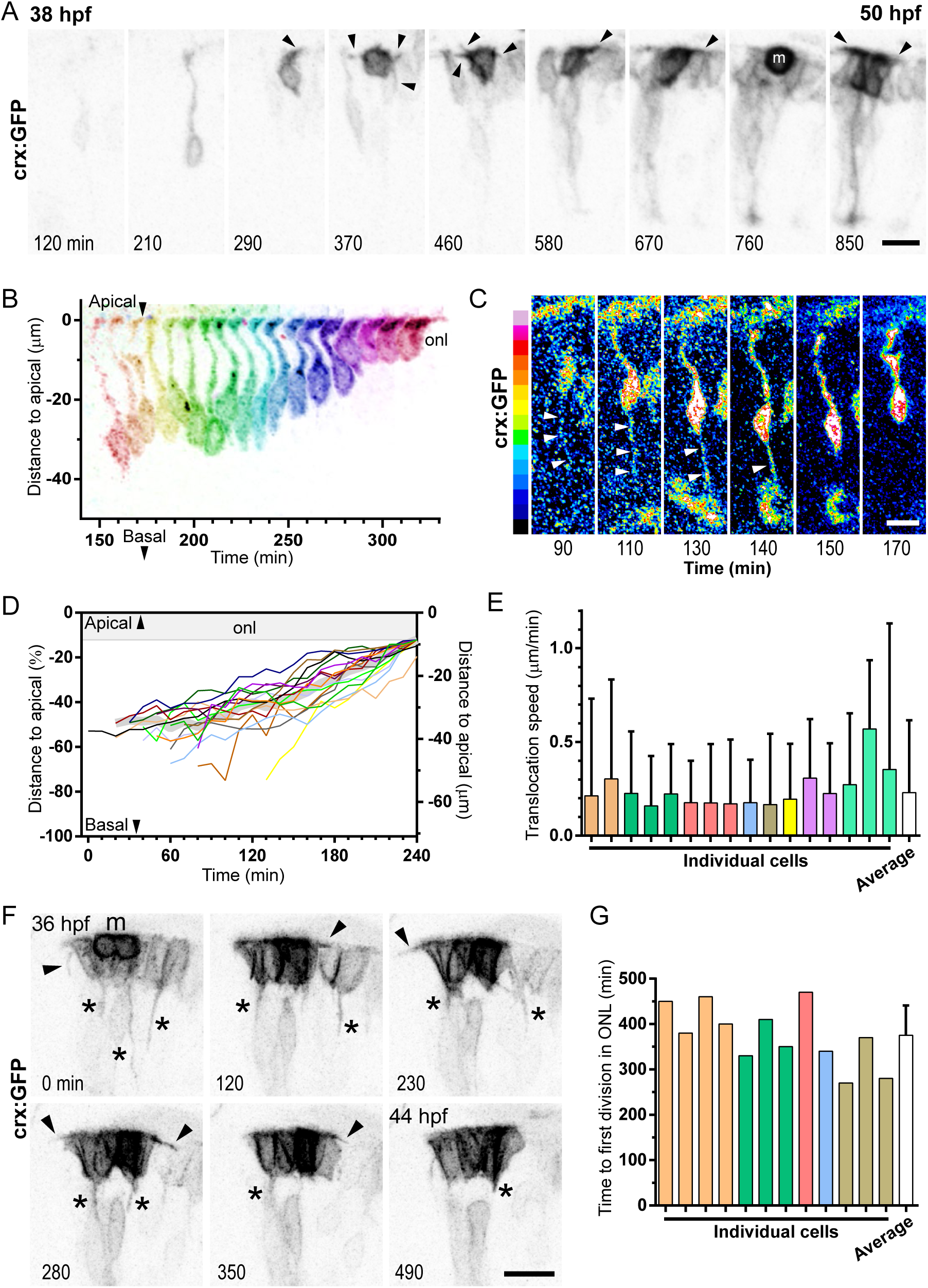
Dynamics of photoreceptor progenitor translocation to the forming outer nuclear layer. **(A)** Maximum intensity projections from a time lapse confocal image of a *crx*:GFP-injected embryo retina, from 36 hpf. A photoreceptor progenitor starts to be detected at time=120 min, and as the *crx*:GFP signal increases (image brightness was adjusted from 580 min to improve visualization), its cell body translocates to the ONL, where it remains, undergoing cell division at t=760 min (cell marked “m”). Arrowheads: cell processes from the apical side of the cell. See Supplementary Video 1. **(B)** Digital superposition of the image of the photoreceptor progenitor observed in A, including all time points (acquired every 10 min) from t=150 to 320 min. **(C)** Observation of the same cell at earlier stages, increasing brightness and converting to a 16-color intensity code, to evidence the presence of a thin basal process (arrowheads) while the cell body translocates basally. Basal detachment and complete retraction of the basal process appears to happen between t=140 and 150 min, just before the cell starts to move apically. See Supplementary Video 2. **(D)** Trajectories in time of several *crx*:GFP-positive cell bodies, since they are detected until they reach the ONL (“onl”). Distance from the base of the cell body to the apical surface was measured for every cell, at each time point, and distances normalized to the width of the neuroepithelium (70.8 µm, represented on the right y axis). Two of the cells did not reach the final position at the end of the time lapse (depicted orange and black). The thick gray line at the back shows the average trajectories of all cells, excluding these two. n=16 cells, 8 embryos, 3 independent experiments. **(E)** Averaged instantaneous speeds of nuclear translocation for each cell shown in D (calculated between the 10 min-separated time points); bars show mean + SD, and colors represent cells from different embryos. The white bar shows the average instantaneous speeds for all cells, and SD (0.23 ± 0.39 µm/min). **(F)** Dynamics of a group of *crx*:GFP labeled photoreceptor progenitors, observed by time lapse confocal microscopy (maximum intensity projections). Arrowheads: cell processes from the apical side of the cell; asterisks: basal processes; time in min. See Supplementary Video 3. **(G)** Time between apical positioning at the ONL and first cell division of 12 different *crx*:GFP-positive cells (6 embryos; 3 experiments). White bar: mean + SD. Scale bars: 10 µm.

**Figure 3.**
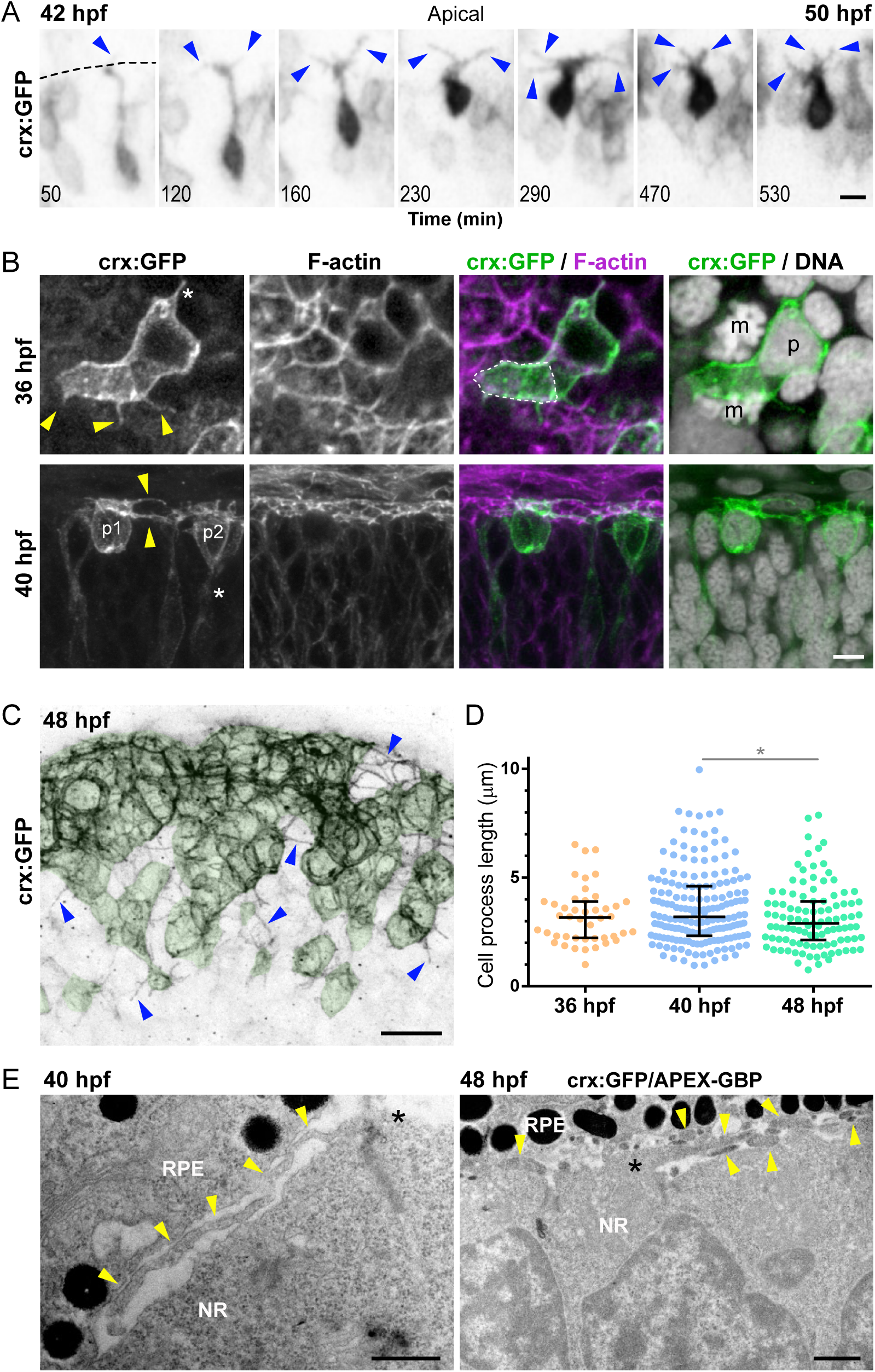
Photoreceptor progenitors extend dynamic cell processes while forming the outer nuclear layer. **(A)** Maximum intensity projections from a time lapse confocal image of a *crx*:GFP-injected embryo retina. An isolated photoreceptor progenitor dynamically extends and retracts tangential processes (blue arrowheads) from 42 to 50 hpf. Dashed line: apical region. **(B)** Confocal images of *crx*:GFP transgenic embryos, tangential (36 hpf) and perpendicular (40 hpf) to the apical retina. Yellow arrowheads: tangential processes; asterisks: basal processes; m: mitotic cell; p: photoreceptor progenitor nucleus; white dashed line: apical membrane of a photoreceptor progenitor. Maximum intensity projections of 4 (36 hpf) and 14 (40 hpf) sections, at a 0.37 µm separation. See Supplementary Videos 5 and 7. **(C)** Tangential view of a 48 hpf retina, at the periphery of the differentiating nasal-ventral patch (maximum intensity projection of 14 sections, at 0.5 µm separation). Blue arrowheads: tangential processes; green shaded area: photoreceptor progenitor cell bodies. **(D)** Comparative quantification of tangential process length at different stages (median and 25/75% percentile). N: 36 hpf, 44 processes from 9 embryos; 40 hpf, 174 processes, 12 embryos; 48 hpf, 103 processes, 9 embryos. Statistical significance analyzed using Mann-Whitney nonparametric test; 40-48 hpf p=0.043). **(E)** TEM images showing the apical portion of the neural retina and the sub-retinal space, where tangential processes are observed (yellow arrowheads). At 48 hpf, processes appear electron-dense because of APEX-GBP detection on *crx*:GFP-positive cells. Scale bars: A, 3 µm; B, 5 µm; C, 10 µm; E, 500 nm

During this period, and particularly between 36 and 48 hpf, many photoreceptor precursors exhibited a high cell cortex activity, with relatively long, thin processes mostly extending parallel to the apical retinal surface, with some arising from basal-lateral regions (Fig. 2A,B,F, Fig. 3A and Supplementary Videos 1, 3, 4 and 5). The highly dynamic nature of these processes can be easily appreciated in Supplementary videos 4 (early-stages *atoh7*:GFP labeling) and 5 (*crx*:GFP). In the anterior-ventral area of 36 hpf retinas, as well as at the periphery of the differentiating cell patch at later stages, isolated *crx*:GFP-positive cells appeared separated from others by a relatively short distance. Some of these cells displayed relatively irregular shapes, with prominent cell protrusions along the apical surface (Fig. 3B and Supplementary Videos 6 and 7). A bit later, at 40 hpf, it was common to see thin, and sometimes branched, processes extending very near and parallel to the apical surface, apparently connecting labeled cells (from here called “tangential processes”; Fig. 3B and Supplementary Video 8). At 48 hpf, a similar behavior was observed in the youngest progenitors, around the periphery of the *crx*:GFP-positive patch of cells (Fig. 3C), but not at 60 hpf. These highly dynamic processes were apparently more numerous at 40 (5-7 simultaneously per cell) than at 36 and 48 hpf (3-4 per cell), but presented similar average lengths of around 3 μm along these stages (Fig. 3D). Ultrastructural observations showed that these tangential processes were extremely thin (around 0.2 µm in diameter), and extended from the apical membrane of *crx*:GFP-expressing cells into the sub-retinal space (Fig. 3E). Occasionally, the tangential processes were observed to closely contact other neural retina apical cells, as well as RPE cells. Remarkably, at 48 hpf, the sub-retinal space appeared to contain many processes crossing the retinal apical surface in all directions (Fig. 3E), something that had been suggested by the apical crisscrossed *crx*:GFP pattern observed by confocal microscopy (see for example Fig. 3C).

### Presence and role of primary cilia in photoreceptor progenitors

Bearing in mind that the connecting cilium is essential for the extension of the photoreceptor outer segment post-mitotically, and that primary cilia are necessary in proliferating retinal progenitors to ensure the correct balance between RGCs and photoreceptors (Lepanto, Davison, *et al.*, 2016), we wondered if *crx*:GFP-expressing photoreceptor precursors would have a primary cilium, and if there is a particular cellular localization and a functional role for this organelle during the formation of the ONL. The transgenic labeling by the expression of Arl13b-GFP (Borovina *et al.*, 2010) together with an immunodetection of γ-tubulin at 40 hpf, revealed the presence of several cilia at the outer retinal surface (Fig. 4A). By labeling cilia using an anti-acetylated tubulin antibody on *crx*:GFP transgenic embryos, we found that, in all observed cases, they were present at an apical localization in photoreceptor progenitors at 36 hpf, and that this localization was maintained through different stages up to 72 hpf (Fig. 4B). The density of cilia at the apical border of the neuroepithelium was maintained (or slightly increased) between 36 and 40 hpf, being that at the former most of the cilia corresponded to non-specified neuroepithelial cells and at the latter, to photoreceptor progenitors (Fig. 4C). Beyond 40 hpf, this density decreased steadily until 60 hpf, with a new apparent increase by 72 hpf (Fig. 4C). The ultrastructural analysis demonstrated that at all these stages, cilia observed on photoreceptor precursors were directed towards the sub-retinal space and had the classical conformation of primary cilia (Fig. 5A). Contrary to what we previously described for neuroepithelial cells and RGC progenitors (Lepanto, Davison, *et al.*, 2016), most cilia found on putative photoreceptor progenitors between 40 and 60 hpf sat on a relatively flat area of the apical membrane, with no evident ciliary pocket (Fig. 5A). In addition to becoming scarcer, cilia showed a tendency to turn shorter along this developmental period (Fig. 5A,C). At 72 hpf, when all photoreceptors are already post-mitotic and starting to grow their outer segments, cilia length appeared to increase again (Fig. 5A,C). These late cilia, however, looked different in shape, being wider in their central-distal portion and usually presenting a partial or complete ciliary pocket (Fig. 5A,C). In spite of these conspicuous changes in length and general shape, two parameters were maintained apparently constant: the diameter of cilia at their base, and that of the electron-dense area of the periciliary plasma membrane (Fig. 5D,E).

**Figure 4.**
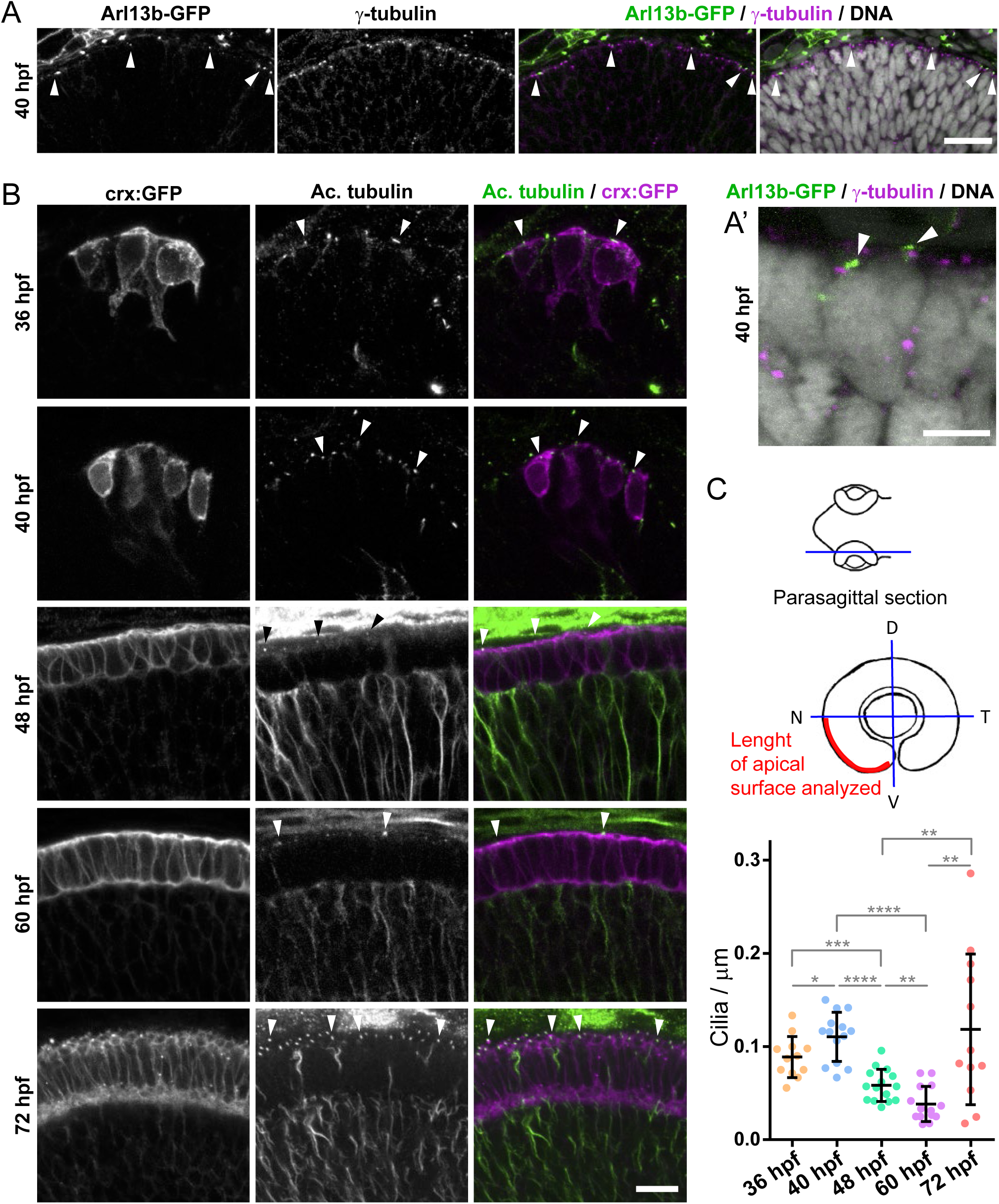
Presence of primary cilia on photoreceptor progenitors. **(A)** Confocal image (maximum intensity projection of 2 sections, 1 µm separation) of the apical region of a 40 hpf zebrafish retina, where cilia are transgenically labeled by Arl13b-GFP (arrowheads) and centrosomes/basal bodies with an antibody to γ-tubulin. In A’, a higher magnification from another section is shown, where two short cilia associated with basal bodies are clearly localized at the apical portion of putative photoreceptor progenitors. **(B)** A similar signal is observed by immunolabeling cilia with an anti-acetylated tubulin antibody on *crx*:GFP transgenic embryos. Very short cilia can be detected at the apical side of photoreceptor progenitors at all stages observed (arrowheads). **(C)** Quantification of the number of cilia along a line spanning the nasal-ventral quarter of the retina on single confocal sections. Mean ± SD are represented by lines. Statistical significance was determined using the Student’s *t* test. Scale bars: A, 20 µm; B, 10 µm.

**Figure 5.**
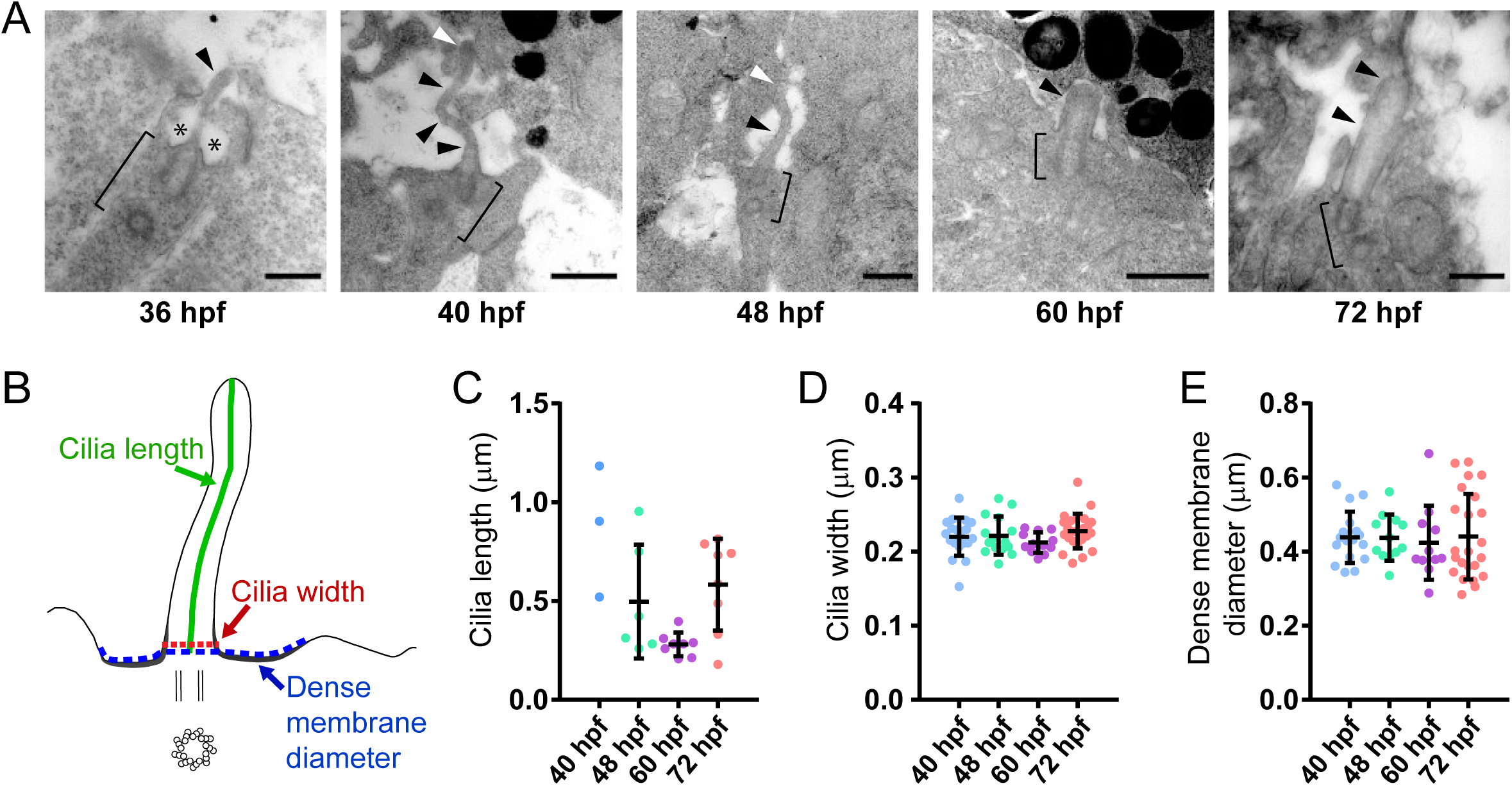
Ultrastructural characterization of primary cilia on photoreceptor progenitors. **(A)** TEM images of ultrathin sections from the apical region of zebrafish retinas at different developmental stages, showing typical primary cilia (arrowheads), all of which are on the apical cell membrane and directed towards the sub-retinal space. At 36 hpf, the cell could not be identified, and the cilium shown probably belongs to a neuroepithelial cell. This cilium presents a deep ciliary pocket (asterisks). Between 40 and 60 hpf, most cilia present on putative photoreceptor progenitors are characterized by the absence of a ciliary pocket, and only very few and extremely short cilia could be detected at 60 hpf. At 72 hpf, photoreceptor cilia were usually larger than at earlier stages. Brackets: basal bodies. **(B-D)** Quantification of primary cilia length, width (at their base) and diameter of the periciliary electron-dense membrane, measured on ultrathin sections imaged by TEM. The drawing in B graphically illustrates the measured parameters. Mean ± SD. Scale bars: 500 nm.

To test for the possible functions of primary cilia in the early stages of photoreceptor polarization and differentiation, we treated embryos to severely disrupt cilia using a combination of morpholino oligomers directed to block the mRNA splicing of two different proteins involved in ciliogenesis: Elipsa and IFT88 (characterized in Lepanto et al., 2016b). Embryos thus treated presented a general “ciliary phenotype”, characterized by a severe ventral curvature of the whole body, and reduced eyes (Fig. 6A). As previously described by us, the neural retina of treated embryos was smaller and with fewer cells than in controls, with a remarkable general delay in cell differentiation (Lepanto, Davison, *et al.*, 2016). In that work, we specifically concentrated on RGCs, but here we used *crx*:GFP transgenic expression to follow the dynamic behavior of photoreceptor progenitors in cilia-impaired embryos. We found that in morpholino-treated embryos, these cells nevertheless formed a tight layer of cells that appeared to be correctly polarized and localized at the apical-most aspect of the retina (Fig. 6B and Supplementary Video 9). An increase in image brightness, however, revealed that all along the registered period (36-53 hpf approximately), many of the photoreceptor progenitors presented long and thin basal processes, a feature seldom seen in control cells beyond the earlier stages (Fig. 6B and Supplementary Video 9; compare to data shown in Fig. 2). These processes appeared to be detached from the basal surface, and displayed a dynamic behavior characterized by alternating periods of elongation and retraction, as visualized in time-lapse confocal images (Fig. 6C and Supplementary Video 10). In all observations made, they appeared not to extend beyond the apical border of the RGCs layer (Suppl. Fig. 1). Several cells actually presented two or more processes, eventually branched, that alternated their periods of extension and retraction (Fig. 6C and Supplementary Video 10). In both control and morpholino-treated retinas, photoreceptor progenitors stopped cell process extension during cell division (Supplementary Video 11).

**Figure 6.**
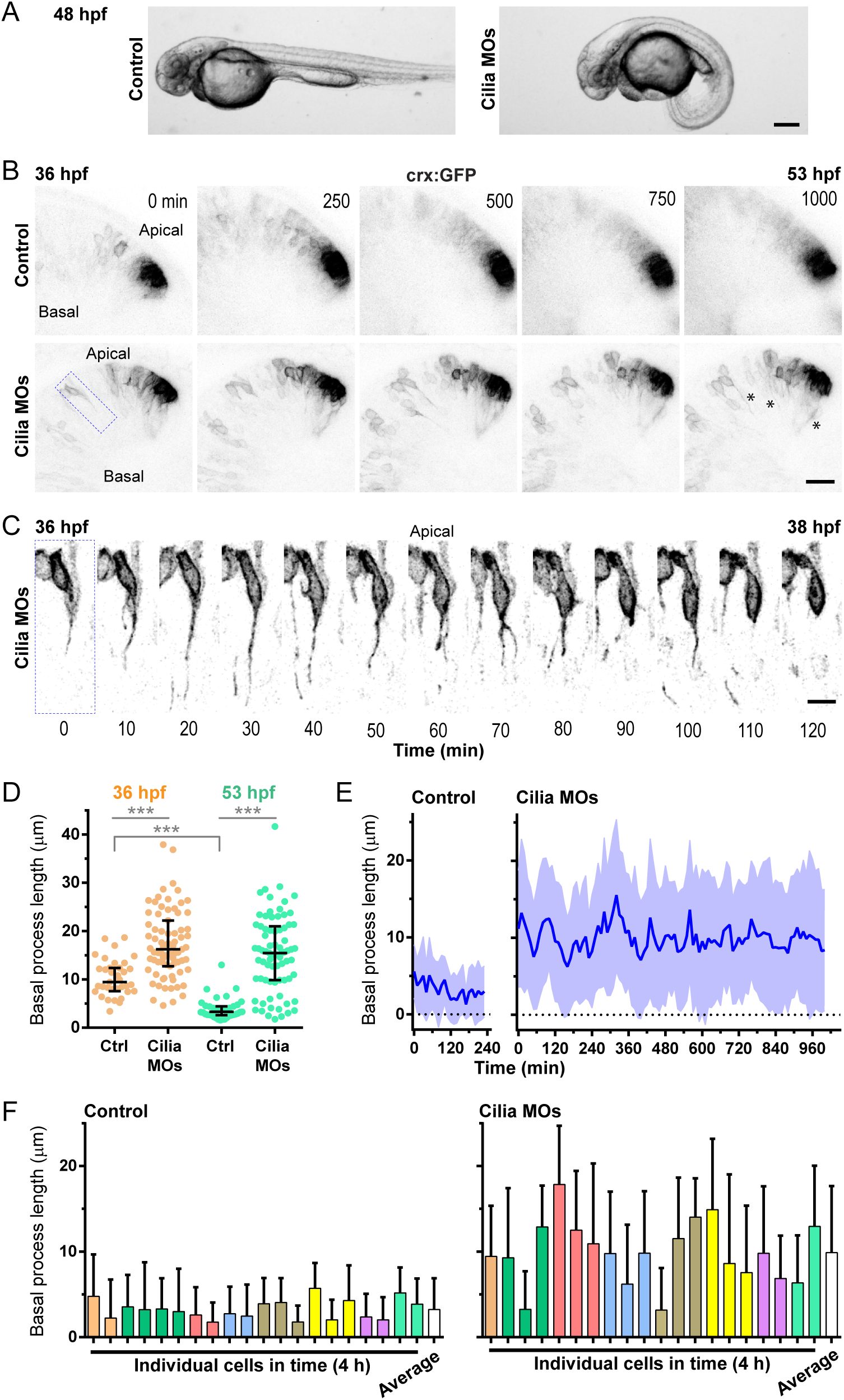
Effect of primary cilia disruption on photoreceptor progenitors. **(A)** Cilia disruption using morpholino oligomers (MOs) to Elipsa and IFT88 caused a clear “cilia defect” phenotype on zebrafish embryos. **(B)** Maximum intensity projection from a time-lapse confocal observation of control and morphant embryos, between 36 and 53 hpf. Long basal processes remain all along the observation time in morphants (asterisks). **(C)** Higher magnification and thinner projection of the cell squared in B, to show all time points in a two-hour period. The basal process is seen to rapidly change in length and shape during this short period. **(D)** There is a significant decrease in basal process length (median ± interquartile values) in control embryos between 36 and 53 hpf, but not in treated embryos (Mann-Whitney test). In both stages, basal processes are significantly longer in morphants. N=5 processes/embryo, from 7 control and 15 morphant embryos at 36 hpf; 8 and 15 embryos, respectively, at 53 hpf (3 independent experiments). **(E)** Averaged (mean ± SD) length of basal processes measured at each time point from individual cells, for 4 hours in controls and 17 hours in morphants. Between 2 and 4 cells/embryo were manually tracked, measuring the length of the basal process on confocal stacks. N=20 cells from 8 embryos; 3 independent experiments in controls, 2 in morphants. **(F)** Analysis of the averaged basal process length (mean + SD) for each cell along the first 4 hours of time-lapse, using the same data as in E. Bars represent individual cells, and each color a different embryo; white bars represent the averaged values. Scale bars: A, 200 µm; B, 20 µm; C, 8 µm.

To further analyze the behavior of these aberrant basal processes, we quantitatively compared their occurrence and dynamics in control and morphant embryos. At 36 hpf, basal processes of variable length were extended from *crx*:CGP-positive in both situations, although they were significantly shorter in controls (Fig. 6D). At 53 hpf, however, very few basally-directed cell processes could be detected arising from the ONL in control embryos, and the few that could be measured were much shorter than the several more observed in cilia-impaired retinas (Fig. 6D). At every time point of time-lapse observations between 36 and 53 hpf, retinas from control and cilia-impaired embryos presented *crx*:GFP-positive basal processes of varying length. In the case of controls, individual basal processes could only be reliably followed for approximately the first four hours (up to 40 hpf), which is graphically represented in Fig. 6E. Here, the average process length at 36 hpf was around 5 µm, while by 40 hpf it was around 3 µm, showing a tendency to becoming shorter as development of the outer nuclear layer proceeded. In treated embryos, on the other hand, we could follow several processes until the end of the recorded period (53 hpf). These processes were much longer in average (around 10 µm), and with a greater variation. Both average and standard deviation were maintained along the 17 hours of time-lapse in morpholino-treated embryos (Fig. 6E). In addition, the length of the basal processes in each cell varied prominently along time in both control and morpholino-treated situations, although both the average length and standard deviation was again much higher in the latter (Fig. 6F).

### The role of N-cadherin in the organization of the outer nuclear layer and early photoreceptor differentiation

With the aim of starting to understand the possible functions of the adhesion molecule N-cadherin in the epithelial-like organization of photoreceptor progenitors in the zebrafish neural retina, we knocked its expression down using extensively characterized morpholino oligomers that have been shown to largely phenocopy the null mutants (Lele *et al.*, 2002) (Fig. 7A). To have a more complete assessment of cell behavior in the neurogenic retina, we performed these knock-downs on double transgenic *crx*:GFP/*atoh7*:gap-RFP (*atoh7*:RFP) embryos (Fig. 7B). This double labeling allowed us to visualize and differentiate the photoreceptor cell layer from the other cell types in the retina, especially other early-born neurons such as retinal ganglion, amacrine and horizontal cells. In addition, bipolar cells, which are late-born neurons also expressing Crx, are not labeled by *atoh7*:RFP and can thus be differentiated from photoreceptor progenitors. At 72 hpf, we observed the expected general disruption of retinal tissue, with the appearance of both ectopic *atoh7*:RFP and *crx*:GFP signal (Fig. 7B). Interestingly, the strongest signal in each case (RGCs and photoreceptors, respectively), appeared to be completely segregated. Similar observations could be obtained after labeling *crx*:GFP embryos for F-actin, which is normally accumulated at plexiform layers at this stage (Fig. 7C). In morphant embryos, most photoreceptors were grouped in small round structures, with a rosette-like appearance. To have a quantitative idea of photoreceptor misplacement, we measured the distance from the cell nucleus to the apical surface in 72 hpf retinas (Fig. 7D). While in controls photoreceptors were confined to the outer nuclear layer, within 14-15 µm from the apical surface, in N-cadherin morpholino-treated embryos we found an enormous dispersion of the cells across the apico-basal axis. In this case, however, photoreceptor distribution presented a preference for the central one third of the retina: 50% of the *crx*:GFP-positive photoreceptors were localized between 40.6 and 68.3% of the retinal width in morphants, against 8-12.4% in controls (Fig. 7D).

**Figure 7.**
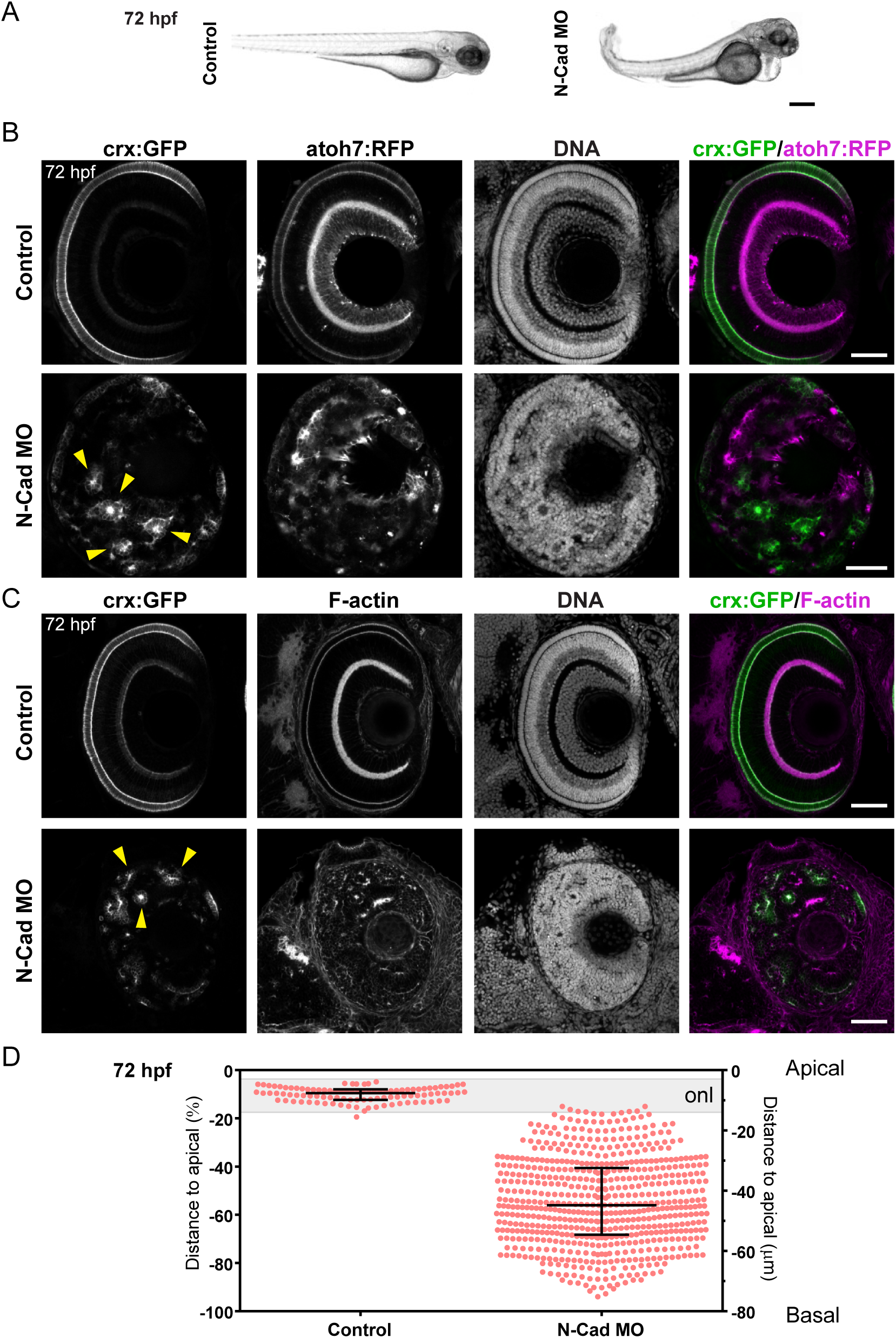
Effect of N-cadherin knock-down on the formation of the outer nuclear layer. **(A)** External phenotype of an N-cadherin morphant (N-Cad) at 72 hpf, compared with a control embryo. **(B)** Single confocal sections of the retina from 72 hpf *crx*:GFP/*atoh7*:RFP double transgenic zebrafish embryos injected with the N-Cad or control MOs, labeled with methyl green (DNA). A severe disruption in retinal layering is observed in morphant embryos, characterized by the formation of multiple photoreceptor-only rosettes (arrowheads). **(C)** Single confocal sections of the retina from 72 hpf *crx*:GFP transgenic embryos injected with the N-Cad or control MOs, labeled with TRITC-phalloidin (F-actin) and methyl green (DNA). **(D)** Representation of the position of photoreceptors along the apico-basal axis of the retina, in control and N-Cad morphants at 72 hpf. The distance of the cell nucleus from the apical surface (position “0”) was normalized as a percent of total retinal width. Median ± interquartile values are shown. N=100 control and 629 morphant cells; from 20 control and 30 morphant embryos; 5 independent experiments. The scale on the right shows the average width of the retina in controls (80 µm), which was indistinguishable from that of morphants (median of 80.4 and 81.2 µm, with SD of 11.5 and 10.5 µm, respectively). The gray band represents the approximate position of the ONL (“onl”) in control embryos. Scale bars: A, 300 µm; B and C, 50 µm.

At higher magnification, the rosette organization of photoreceptors in N-cadherin morphants was more evident (Fig. 8A,B). In these structures, photoreceptors appeared wedge-shaped, with a smaller portion towards the center of the rosette, and a wider side towards the periphery. Both F-actin and the polarity marker aPKCζ were accumulated at the central portion, indicating an apical identity of this region (Fig. 8A,B). The difference in cell shape between control and morphant photoreceptors was partly quantifiable by measuring the cell length between their basal and apical ends (“cell height”, Fig. 8C), where morphant cells were found to be significantly shorter than their control counterparts (median was 13.9 µm in controls, against 6.9 µm in N-cadherin morphants; SD: 1.1 and 1.2 µm, respectively). To further analyze the process that leads to this ectopic position and aberrant organization of the photoreceptor progenitors in N-cadherin-impaired retinas, we followed the behavior of these cells since 36 hpf on *crx*:GFP/*atoh7*:RFP double transgenic embryos by time-lapse confocal microscopy (Fig. 9). At these earlier stages, as differentiation is only evident at the anterior-ventral portion of the retina, this is where structural disruption was also evident in N-cadherin morphants, as a wide extension of the retinal tissue beyond the apical border (Fig. 9A and Supplementary Figure 2). Photoreceptor progenitors could be unambiguously detected from very early stages because they were doubly labeled by GFP and RFP. They were usually first detected in central regions of the retina and then started to display random short movements, eventually leading to some displacement, showing a high degree of cell cortex activity as seen in control photoreceptor progenitors (Fig. 9B,C; Supplementary Figure 2, and Videos 12-15). We thus wondered how photoreceptor rosettes are formed in the N-cadherin morphants. Figure 9B, and Supplementary Video 12, show how two initially isolated *crx*:GFP-positive cells approach and attach, being joined by other cells to form an identifiable rosette in approximately 12 hours. Interestingly, both cells divided at the same time, around 5-6 hours after binding, which is comparable to the quantified time of the first cell division of *crx*:GFP-positive progenitors after arriving to the ONL in control embryos (see Fig. 2G). Supplementary Figure 2A and Video 14 illustrate another example of the process of rosette formation. As photoreceptor progenitors displaced in the retina, they usually extended profuse neurite-like cell processes, similar to the tangential processes we observed in control embryos (Fig. 9E and Supplementary Fig. 2B; Supplementary Videos 13 and 15). A quantification of the length of these processes found in many ectopic *crx*:GFP-positive cells, either isolated or in rosettes, indicated that they are slightly but significantly longer than the tangential processes measured in controls at 40 hpf (Fig. 9F).

**Figure 8.**
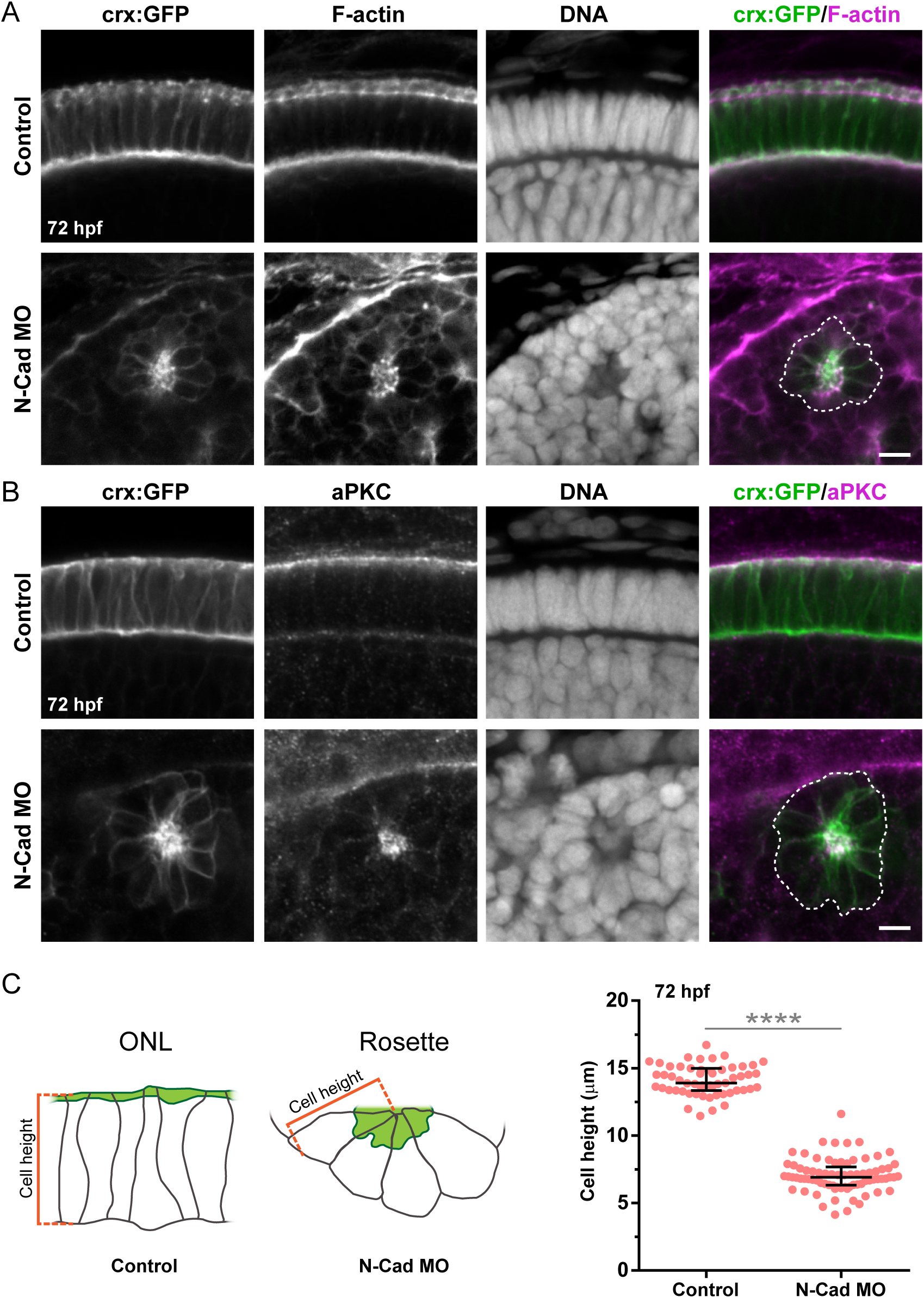
Effect of N-cadherin knock-down on photoreceptor morphogenesis. **(A-B)** High magnification confocal sections of the retina from control and N-Cad MO-injected *crx*:GFP transgenic embryos at 72 hpf, labeled with TRITC-phalloidin (F-actin; A), an anti-aPKCζ antibody (aPKC; B) and methyl green (DNA). Rosette-like groups of photoreceptors (demarcated with the white dashed line) are observed, where both F-actin and aPKC accumulate at the center. **(C)** Comparative quantification of the height (apico-basal length) of photoreceptors in control and N-Cad morphant embryos at 72 hpf. N=55 control and 77 morphant cells; 10 control and 14 morphant embryos; 4 independent experiments. Median and interquartile range are shown; statistical significance was determined using the Mann-Whitney test. Scale bars: 5 µm.

**Figure 9.**
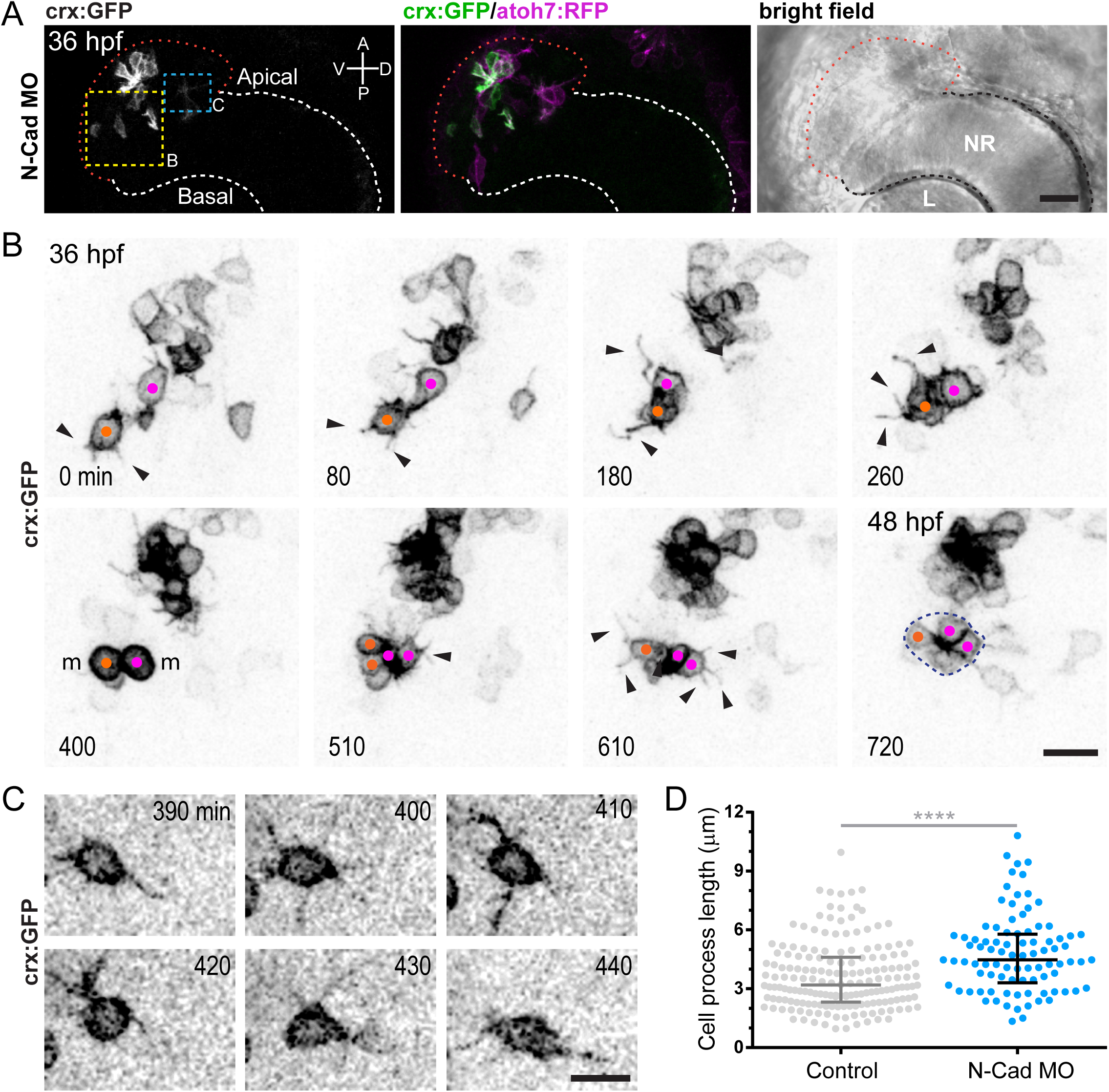
Effect of N-cadherin knock-down on the dynamics of photoreceptor progenitors. **(A)** Low magnification confocal images of the beginning of the time-lapse analyzed in B and C, at the level of the anterior region of an N-cadherin morphant retina, from a *crx*:GFP/*atoh7*:RFP double transgenic embryo. L: lens; NR: neural retina. White dashed line: limit of the neural retina; red dashed line: apical limit of the aberrant anterior-ventral area of the retina. **(B)** Detail of confocal time-lapse experiment showing the process of rosette formation by ectopic photoreceptor progenitors (yellow dashed area in A). The color dots indicate individual cells that join to form the rosette, and their respective daughter cells. These cells mitotically divide at the same time (m). Arrowheads: cell processes. **(C)** Magnified image of another region of the same time-lapse (blue dashed area in A), showing a single *crx*:GFP/*atoh7*:RFP-positive cell at different time points, to better visualize the neurite-like processes. **(D)** Quantification of photoreceptor progenitors cell process length in N-cadherin morphants at 40 hpf, compared with the length of tangential processes in normal embryos (control data are the same as in Fig. 3D). N-cadherin morphants N=89 processes; 8 embryos; 3 independent experiments. Median and interquartile range are indicated; statistical significance determined using the Mann-Whitney test. Scale bars: A, 20 µm; B and C, 10 µm.

Finally, in order to better characterize the behavior of photoreceptor progenitors in N-cadherin knock-down retinas, we tracked the trajectories of these cells from the moment they were first detected by *crx*:GFP expression to the inclusion in a rosette. Even if they did not translocate for long distances, the observed cells displayed apparently random movements, with different speeds and directions (Fig. 10A). We also compared the time it took the morphant photoreceptors to reach a rosette, to that of controls to translocate to the ONL, finding it was significantly longer (nearly double: 346 ± 70 against 189 ± 31 min, respectively) (Fig. 10B, and see also Fig. 2 for more data on untreated embryos). The effective displacement (Euclidean distance between the starting and destination points) was in all cases much shorter than the total distance migrated (Fig. 10C), which is strong indicator of a random movement. Also using the data from the tracked trajectories, we could determine the instantaneous and average speeds of these cells while migrating (Fig. 10D,E). Similarly to what we describe above for apical translocation in normal retinas, there was a high level of variability in cell speed at different time points (Fig. 10D, compare to Fig. 2E), and somehow surprisingly, the average migration linear speed in N-cadherin-treated embryos was statistically undistinguishable from the translocation linear speed in normal embryos, with median values around 0.3 µm/min (Fig. 10E).

**Figure 10.**
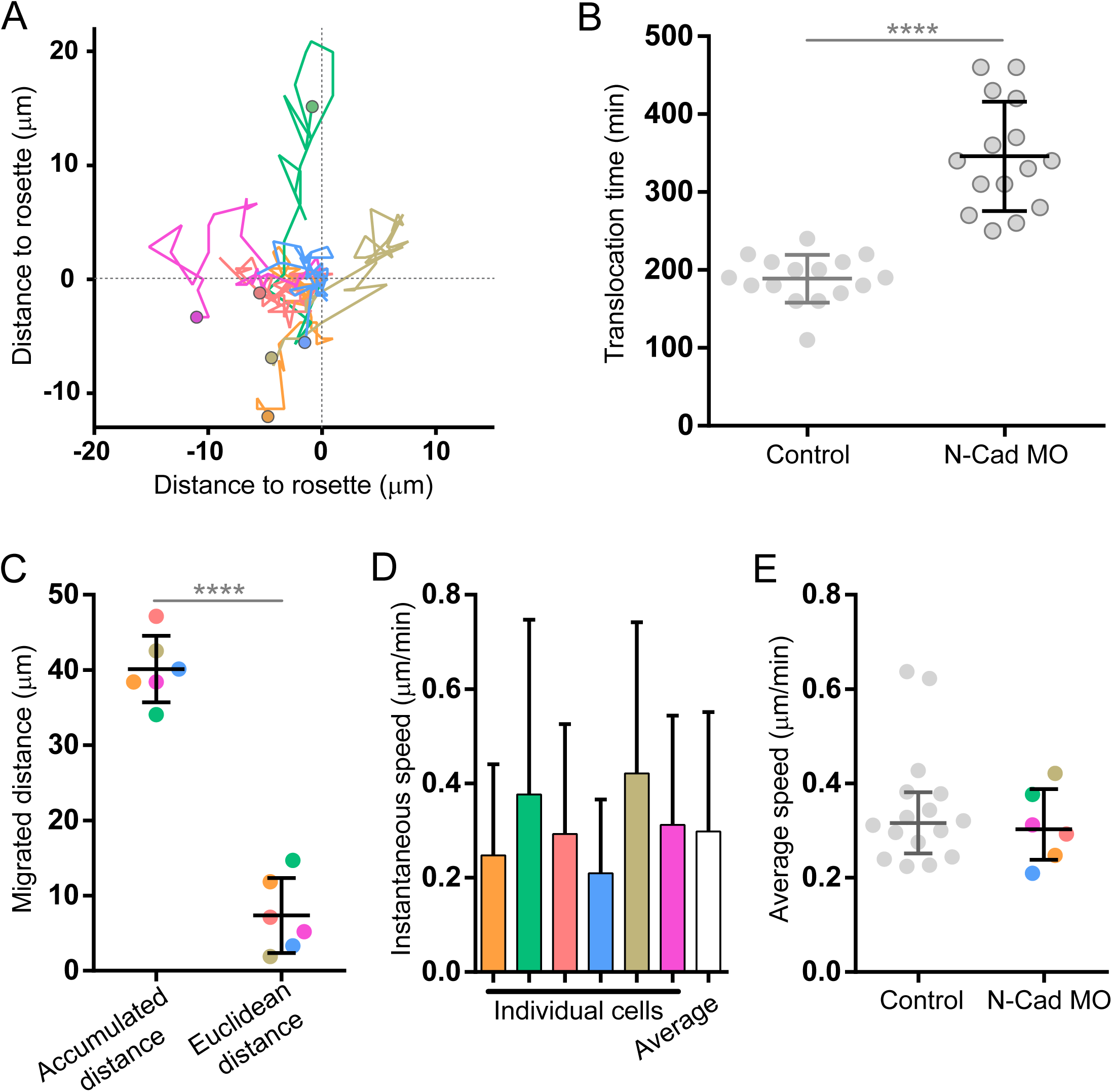
Quantitative analysis of ectopic photoreceptor progenitor dynamics in N-cadherin morphants. **(A)** Tracked trajectories (x-y) of 6 *crx*:GFP-positive cells (3 embryos, 2 independent time-lapse experiments), from the onset of GFP detection to their positioning at a rosette. All trajectories are aligned to the final position (0 µm in both axes) and the starting point is marked with a colored dot. **(B)** Comparison of the time that *crx*:GFP-positive cells take to become inserted in the ONL (Control) or in a rosette (N-Cad MO), since the time GFP signal becomes detectable. Control data are from the same data set analyzed in Fig. 2D-E, and N-Cad MO data are the same as A in this figure. Mean ± SD; statistical significance determined using the Student’s *t* test. **(C)** Comparison of the total distance migrated by photoreceptor progenitors (Accumulated distance) with the effective displacement (Euclidean distance) in N-cadherin morpholino-treated embryos. The data used for this calculation were the same as in A, maintaining the color code for cell identification. Mean ± SD; statistical significance determined using the Student’s *t* test. **(D)** Instantaneous speeds of photoreceptor progenitors from A (same color code), averaged along the whole time. White bar: averaged instantaneous speeds of all cells. Mean + SD. **(E)** Comparison of the average speed of migration in N-cadherin morphants (N-Cad MO) to the translocation speed in normal (Control) embryos, as shown in Fig. 2E. Median ± interquartile range; medians no statistically different according to Mann-Whitney test.

## DISCUSSION

Previous research on vertebrate photoreceptors differentiation has largely concentrated either on the initial cell fate decisions that give place to the generation of committed progenitors (see for example Boije *et al.*, 2014), or on the maturation and differentiation of the outer segment (Wheway *et al.*, 2014). In the present work, we decided to start filling the gap between these two processes that are not only apart in terms of cell differentiation stages, but also in time, since one of the most salient particularities of differentiating photoreceptors is the long time they spend between specification (highlighted by the expression of distinctive markers such as Crx; Shen and Raymond, 2004) and actual cell differentiation (Crespo and Knust, 2018). Remarkably, cones are among the first cell types to be specified in the retina, even sharing the expression of the proneural transcription factor Atoh7 with the first neurons to become post-mitotic, RGCs (Boije *et al.*, 2014). RGCs start to differentiate a very short time after the last cell division at the apical side of the neuroepithelium. The stereotyped differentiation process of these cells occurs in a few hours in the zebrafish, and it does not include an “unpolarized” stage, but a gradual transition in which the nascent neuronal polarity overlaps with the fading epithelial polarity (Zolessi *et al.*, 2006). We have demonstrated here some relevant differences in the way photoreceptors differentiate: first, cycling photoreceptor progenitors translocate their nuclei and accommodate at the final place of differentiation many hours before becoming posmitotic (our quantification shows 6:25 ± 1:06 h to the first cell division, which happens around 48 hpf, and Weber *et al.*, 2014, showed that the second and last cell division occurs 12-24 hours after this); second, cell body translocation to reach this position is apical-ward, and they detach from the basal lamina to retract a basal process; third, once at the apical side, and albeit showing evident signs of polarization such as an apical cilium and adhesion complex molecules, they spend several hours extending multiple neurite-like cellular processes along the sub-retinal space. These tangential processes, which to our knowledge have not been previously described, are rather remarkable in cells that when mature only have a very short axon and no dendrites, and are reminiscent of the unpolarized stage 2 of rat hippocampal neurons in culture (Cáceres *et al.*, 2012). We have previously shown that RGCs can also display this type of behavior in culture, or *in vivo* when lacking a positional signal for axonogenesis, such as Laminin1 (Randlett *et al.*, 2011). Interestingly, the tangential processes in photoreceptor progenitors appear to peak in number and length around the time when the ONL is being assembled, to stop by around the time photoreceptors undergo their last cell division and start to elongate their bodies after 60 hpf. It is tempting to speculate that they might have a function in establishing contacts between early photoreceptors, helping in migration and leading to their organization in the crowded and highly regular ONL. In support of this supposition, we showed here that isolated ectopic photoreceptor progenitors in N-cadherin morphants display longer neurite-like processes before and during the establishment of rosettes.

Motivated by the observation by us and others (see for example Crespo and Knust, 2018) of an early, pre-neurogenic, epithelial-like polarity of photoreceptor precursor cells, we decided to explore the possible roles of two conspicuously polarized structures in the retinal neuroepithelium: the primary cilium and the sub-apical N-cadherin-based adhesion complex. Primary cilia have been shown to have different functions in neuronal differentiation (Lepanto, Badano, *et al.*, 2016). We describe here the presence of relatively short primary cilia, located at the apical membrane of photoreceptor progenitors, which can be differentiated from those of neuroepithelial cells by their apparent lack of a complete ciliary pocket (Lepanto, Davison, *et al.*, 2016). These cilia get shorter and scarcer as development proceeds, to nearly disappear by 60 hpf, clearly indicating that they are not the direct precursors of the outer segment, a structure that starts to form by around 72 hpf (Branchek and Bremiller, 1984). Early primary cilia did not appear to be essential for the orientation of photoreceptor progenitors or the formation of an ONL, as seen upon their disruption using a previously characterized combination of morpholinos to ciliogenesis-involved proteins. These morpholinos, acting together, cause the knock-down of two independent intraflagellar transport proteins, IFT88 and Elipsa giving a very reliable and reproducible phenotype, with no evident off-target effects (Lepanto, Davison, *et al.*, 2016). Similar to what was described in that previous work, this treatment generated a severe growth and cell differentiation delay in the retina, where all cell types are reduced in number, but maintaining the laminar organization. By using the *crx*:GFP reporter, we could also describe here a very remarkable effect of cilia disruption on the process of basal processes retraction in photoreceptor progenitors. Instead of a very quick retraction after detachment observed in the untreated condition, in morphants these basal processes remained elongated for several hours, even if photoreceptor cell bodies were already positioned and divided normally at the apical retina. An analog phenotype was observed in Slit1b morphants on post-mitotic RGCs, which would take longer than usual to detach and retract the apical process, even if differentiating normally at the basal side (Zolessi *et al.*, 2006). One important difference is that the elongated basal processes of photoreceptors were in general detached from the basal surface and extremely dynamic in length. Hence, contrary to what was described for RGCs, where the failure to retract was due to a deficiency in N-cadherin down-regulation mediated by the Slit receptor Robo3 (Wong *et al.*, 2012), the maintenance of photoreceptor basal processes does not seem to depend on a defect in detachment. Rather, cilia disruption seems to be affecting the shortening of the basal process. At this stage, we cannot ascertain if the observed phenotype is caused cell autonomously, by the disruption of cilia in the affected photoreceptor progenitors themselves, or if it is related to a missing signal from the basal retina, where RGCs are generated in lower than normal numbers (Lepanto, Davison, *et al.*, 2016). This latter idea is supported by the observation that, in spite of their dynamic behavior, basal processes never appeared to extend beyond the apical limit of the ganglion cells layer.

The proper differentiation of most central nervous system neurons requires a down-regulation of epithelial polarity, as was shown for RGCs (Zolessi *et al.*, 2006). In neurons, typical epithelial polarity molecules, such as N-cadherin or Pard3 are no longer associated to an apical identity and appear to gain new functions in axon specification and neurite outgrowth (see reviews in Gärtner *et al.*, 2015; Hapak *et al.*, 2018). Photoreceptors in general (either from vertebrates or invertebrates) have the particularity of presenting, simultaneously, polarity features found in neurons and epithelial cells, providing a very interesting opportunity to understand the roles of apical adhesion complexes in neuronal development and evolution. Here, we concentrated in further characterizing the roles of the adhesion protein N-cadherin on the organization of photoreceptor cells in an ordered layer, and in their polarization or orientation. Like it was previously described for mutants, the downregulation of N-cadherin expression caused a general disorganization of the zebrafish retina, with the extensive formation of cell rosettes, where photoreceptor progenitors polarized with their apical region towards the center (Erdmann *et al.*, 2003; Masai *et al.*, 2003; Wei *et al.*, 2006). Our time-lapse analyses demonstrated that at early stages, and similar to what happens in control embryos, photoreceptor progenitors in morphants for N-cadherin arise from Atoh7-expressing cell precursors that individually translocate (in this case, migrate) to eventually join other photoreceptors (in this case, in rosettes instead of the ONL). This translocation took, however, the double of time in morphant than in control embryos, suggesting either a direct role for N-cadherin in this process, or that the integrity of neuroepithelial polarity is an important factor. It was also interesting that even if the general architecture of the retina was completely altered by N-cadherin knock-down, these non-polarized and apparently disoriented isolated progenitors were able to recapitulate the initial stages of ONL formation, albeit in a small round structure (a rosette): they joined together, divided a few hours later and much later differentiated acquiring an apparently normal photoreceptor-like cell polarity (at least in initial stages). N-cadherin appears then to be necessary for photoreceptors to remain attached to the outer limiting membrane through adherens junctions, but not for these cells to recognize each other and eventually join and correctly polarize.

Altogether, the findings described in this work reinforce the idea of a complex set of signaling processes involved in sculpting the final functional morphology of neurons, and that each neuron requires a unique set, exquisitely tailored to its particular shape, position and function. The very special case studied here, the vertebrate photoreceptor, may not represent “canonical” neurons, but exactly because of its characteristic of maintaining a combination of epithelial and neuronal polarity features, it becomes a unique opportunity to begin to understand how neurons arise from epithelia, both in development and in evolution.

## Abbreviations

aPKC: atypical protein kinase C;
Crx: cone-rod homeobox transcription factor;
hpf: hours post-fertilization;
ONL: outer nuclear layer;
RGC: retinal ganglion cell;
TEM: transmission electron microscopy.

## Acknoledgements

We thank Kristen Kwan, Rachel O.L. Wong, Brian Ciruna and William A. Harris for sharing plasmids and/or fish lines; Gabriela Casanova (Unidad de Microscopía Electrónica, Facultad de Ciencias, Udelar), Álvaro Olivera (CURE Rocha, Udelar) and Marcela Díaz / Tabaré De Los Campos (Unidad de Microscopía, Institut Pasteur Montevideo) for technical assistance with microscopy; Gisell González, for her invaluable and continuous support with fish care. This work was partly funded by an ANII-FCE grant to FRZ (1_1_2014_1_4982); SNB PhD fellowship to GA; CAP-Udelar Master’s fellowship to MR; FOCEM-Institut Pasteur de Montevideo Grant (COF 03/11); Programa de Desarrollo de las Ciencias Básicas (PEDECIBA, Uruguay).

## Supplementary Material Legends

**Supplementary Figure 1.**
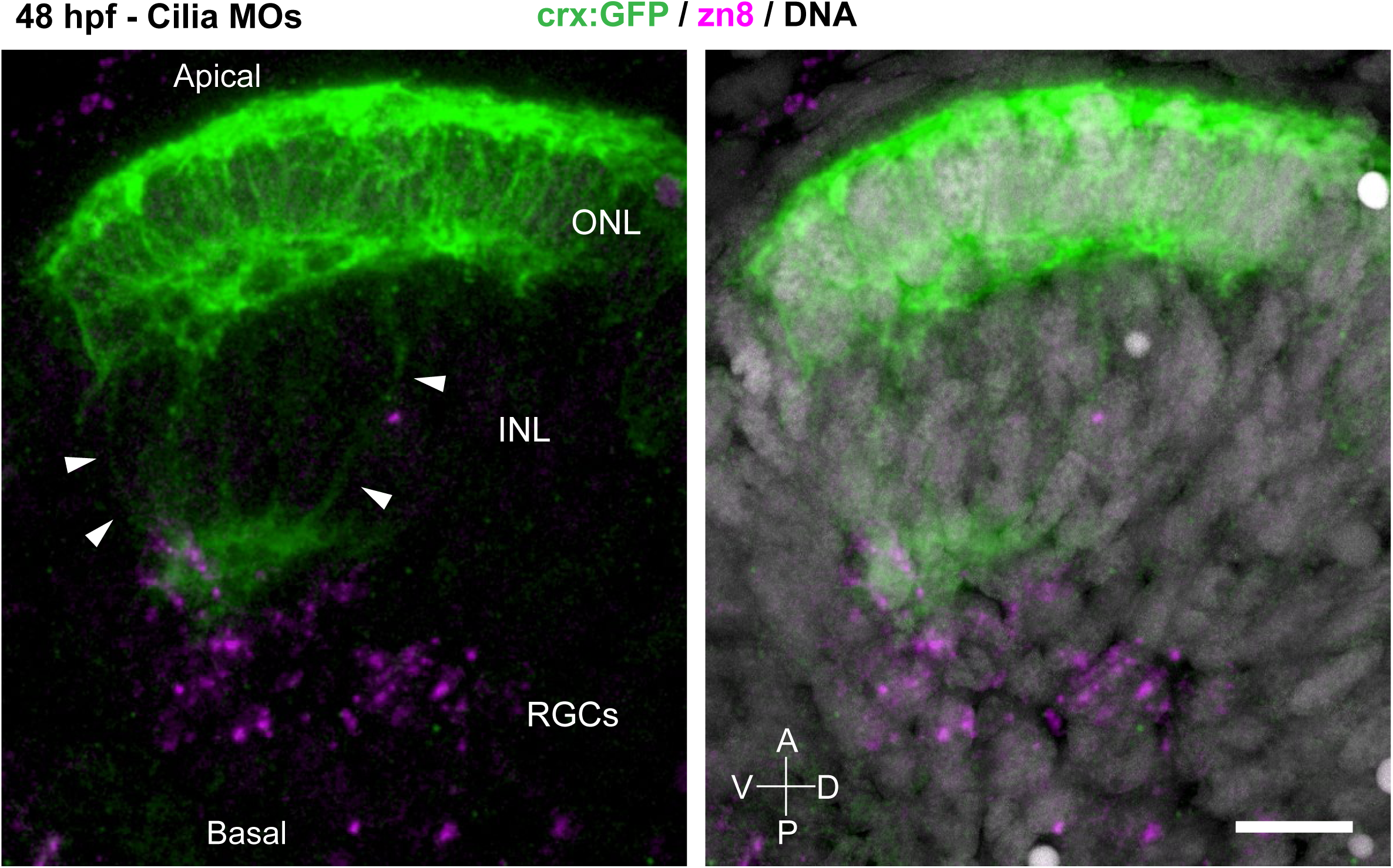
Extension of aberrant basal processes by photoreceptor progenitors in Elipsa/IFT88 morphant retina. Maximum intensity projections of a confocal stack (35 slices, separated by 0.5 µm) showing the anterior-ventral region of the retina from a *crx*:GFP transgenic embryo, labeled to highlight the RGCs (zn8 antibody) and nuclei (methyl green). Arrowheads: basal processes; INL: inner nuclear layer; ONL: outer nuclear layer; RGCs: retinal ganglion cells. Scale bar: 10 µm.

**Supplementary Figure 2.**
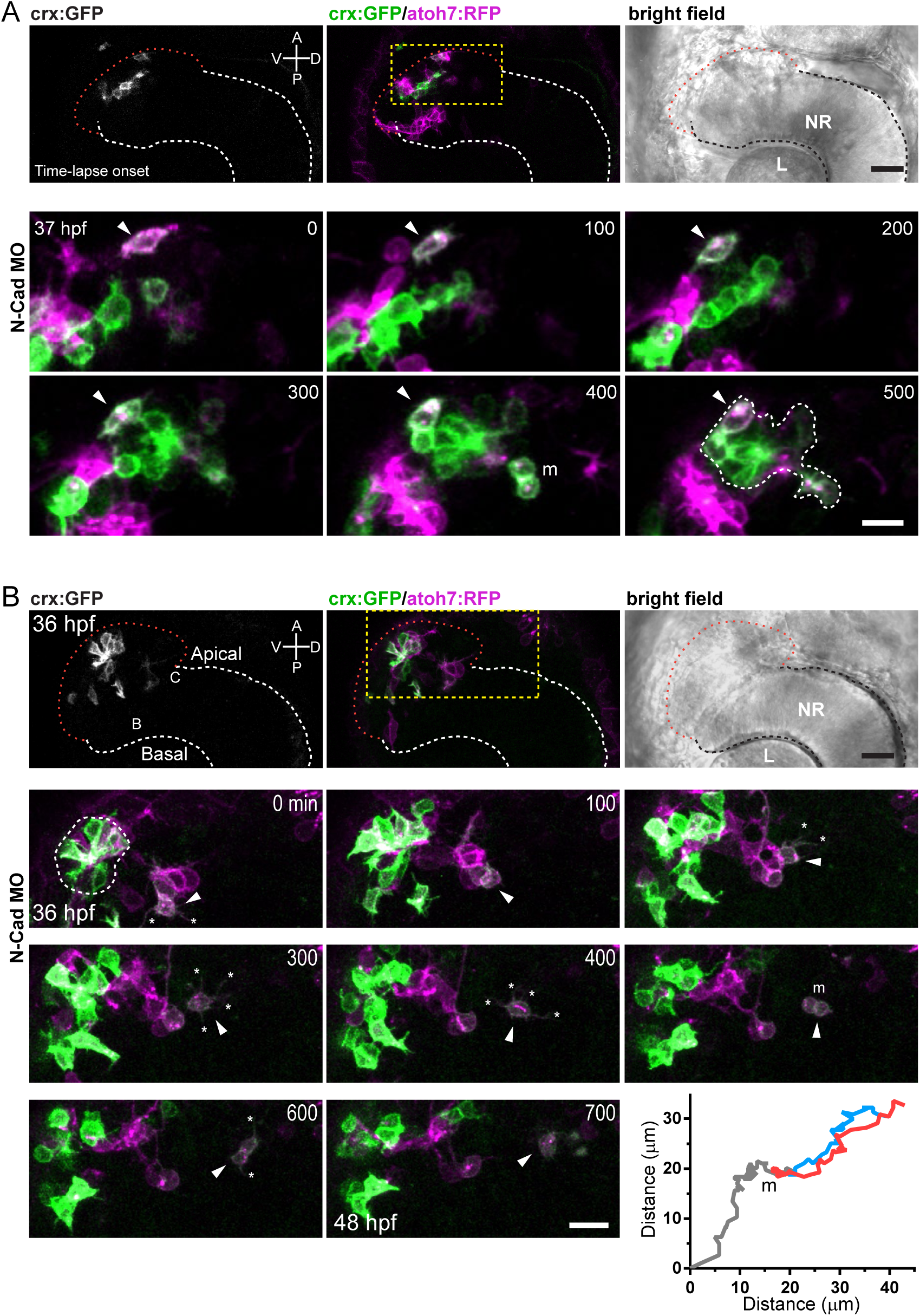
Photoreceptor progenitors dynamics in *crx*:GFP/*atoh7:*RFP double transgenic embryos, upon N-cadherin knock-down: two examples. **(A)** Several disperse cells coalesce to form a single rosette. One of these cells starts expressing high levels of *atoh7*:RFP and to gradually increase *crx*:GFP expression with time (arrowhead). m: photoreceptor progenitor undergoing mitosis, just before the daughter cells join the rosette. **(B)** Dynamics of an isolated photoreceptor progenitor (arrowheads) from the same time-lapse shown in Figure 10. This progenitor is initially detected as a double-labeled cell, around the central retina and in close association with cells strongly expressing only RFP (most probably RGCs). Eventually, the photoreceptor separates from these cells, displaying several dynamic cell processes (asterisks), until it divides at time point 500 min (cell marked “m”). The upper row shows a lower magnification of the embryo eye and head at the beginning of the time-lapse; the yellow squared area corresponds to the time sequence below. A tracking of the cell and its daughters is represented in the graph at the lower right corner. White dashed lines (A-B): limit of the neural retina; red dashed lines: apical limit of the aberrant anterior-ventral area of the retina.

**Supplementary Video 1.** Confocal time-lapse experiment (maximum intensity projection) from a *crx*:EGFP-CAAX (*crx*:GFP) transgenic embryo showing a photoreceptor progenitor since it starts expressing detectable GFP at approximately 38.5 hpf until it integrates to the outer nuclear layer where it divides symmetrically to give rise to two daughter cells that remain there. 3D stacks of confocal sections at 1 µm separation were taken every 10 min for around 17 hours in total. During this time, the cell is seen to detach a basal process (see Supplementary Video 2), translocate its body apically, and display a high cortical activity including the extension and retraction of multiple tangential and basal processes. This video is further analyzed in Fig. 2A-C, as well as in Supplementary Video 2.

**Supplementary Video 2.** Confocal time-lapse experiment (maximum intensity projection of a few sections in each photogram) from a *crx*:GFP transgenic embryo showing the same photoreceptor progenitor as in Supplementary Video 1. A color palette of 16 colors was used to highlight very low fluorescence, like that from the thin basal process that appears to be attached to the basal retinal surface at the initial part of the time-lapse, and detaches between 140 and 150 min. See Fig. 2C for further reference.

**Supplementary Video 3.** Confocal time-lapse experiment (maximum intensity projection) from a *crx*:GFP transgenic embryo showing a group of photoreceptor progenitors already localized at the future ONL. A lateral and a top (from apical) view are shown. These cells continue displaying a high surface activity all along the time-lapse, which includes the extension and retraction of multiple cell processes (mostly tangential and basal). One cell just divided at the beginning of the time-lapse and both daughter cells remain apical. This video is further analyzed in Fig. 2F.

**Supplementary Video 4.** Confocal time-lapse experiment (maximum intensity projection) from an *atoh7*:Gap-EGFP (*atoh7*:GFP) transgenic embryo, showing a cell that can be identified as a photoreceptor progenitor because it remains at an apical position, has no long basal process and progressively reduces GFP expression. This cell continuously extends and retracts many cell processes, including many tangential and a few directed basally.

**Supplementary Video 5.** Confocal time-lapse experiment (maximum intensity projection) from an *crx*:GFP transgenic embryo, showing a photoreceptor progenitor translocating its cell body while profusely extending tangential processes at its apical process tip (see Fig. 3A).

**Supplementary Video 6.** Animation through a confocal stack at high magnification, of a tangential view of a 36 hpf retina from a *crx*:GFP (green) transgenic embryo, labeled with phalloidin (F-actin, magenta) and methyl green (nuclei, blue). A photoreceptor progenitor is seen, which has a prominent apical protrusion squeezed between two mitotic cells. Some short tangential processes extend from this protrusion. See details in Fig. 3B.

**Supplementary Video 7.** Animated 3D reconstruction from a confocal stack at high magnification, of the apical retina from a 36 hpf *crx*:GFP (green) transgenic embryo, labeled with phalloidin (F-actin, magenta) and methyl green (nuclei, blue). A photoreceptor progenitor is seen, which presents a cell protrusion that extends along the retinal apical surface. The cell body of this cell also bulges towards the sub-retinal space, apparently displacing the nuclei of two overlaying RPE cells.

**Supplementary Video 8.** Animated 3D reconstruction from a confocal stack at high magnification, of the apical retina from a 40 hpf *crx*:GFP (green) transgenic embryo, labeled with phalloidin (F-actin, magenta). The photoreceptor progenitor in the center extends many tangential processes radially, some of which appear to contact a neighboring cell, also *crx*:GFP-positive. See details in Fig. 3B.

**Supplementary Video 9.** Confocal time-lapse experiment (maximum intensity projection) showing the retinas from two *crx*:GFP transgenic embryos: control, on the left, and morphant for Elipsa/IFT88 on the right. Recorded time was 17 hours (36-53 hpf), with images taken every 10 min. Although in both cases photoreceptor progenitors accommodate at the apical retina to form an outer nuclear layer, the ones in the morphant present relatively long basal processes that remain highly active all along the time-lapse. See Fig. 6B and C, as well as Supplementary Video 10.

**Supplementary Video 10.** Detail view from the confocal time-lapse experiment (maximum intensity projection) in Supplementary Video 9, showing the behavior of a small group of photoreceptor progenitor in an Elipsa/IFT88 MO-injected embryo. Two cells are particularly evident, displaying very active basal processes that extend and retract. Some other short processes are also evident, either tangential to the retinal surface or extending from the lateral-basal membrane. See Fig. 6C.

**Supplementary Video 11.** Detail view from a confocal time-lapse experiment (maximum intensity projection), from an Elipsa/IFT88 morphant showing a photoreceptor progenitor dividing normally and stopping basal process extension just during M phase.

**Supplementary Video 12.** Confocal time-lapse experiment showing *crx*:GFP-positive photoreceptor progenitors joining to form a rosette. See details in Fig. 9B.

**Supplementary Video 13.** High magnification from the time-lapse image analyzed in Fig. 9 and Supplementary Video 12, showing an isolated *crx*:GFP-positive photoreceptor progenitor extending multiple and highly dynamic neurite-like cell processes. See details in Fig. 9C.

**Supplementary Video 14.** Confocal time-lapse experiment showing *crx*:GFP/*atoh7*:RFP double-labeled photoreceptor progenitors joining to form a rosette (dotted line in last photogram). Asterisks: cell processes. See details in Supplementary Fig. 2A.

**Supplementary Video 15.** Detail view from a confocal time-lapse experiment (maximum intensity projection) from an N-cadherin morphant, showing a double-labeled *atoh7*:RFP/*crx*:GFP cell detaching from a group of *atoh7*:RFP-positive cells to migrate while acquiring a “stage 2-like” conformation with several dynamic neurite-like processes.

